# Behavioral divergence between a generalist and a specialist mosquito despite minimal differentiation across chemosensory receptor gene families

**DOI:** 10.64898/2025.12.07.692824

**Authors:** Marilene M. Ambadiang, Caroline Fouet, Fred A. Ashu, Desiree E. Rios, Aditi Kulkarni, Sourav Roy, Calmes Bouaka, Véronique Penlap-Beng, Colince Kamdem

**Affiliations:** Department of Entomology, Centre for Research in Infectious Diseases, Yaoundé, Cameroon; Department of Biochemistry, Faculty of Science, University of Yaoundé 1, Yaoundé, Cameroon; Department of Biological Sciences, The University of Texas at El Paso, El Paso, Texas, USA; Center for Global Infectious Diseases Research, Seattle Children’s Research Institute, Seattle, Washington, USA; Department of Vector Biology, Liverpool School of Tropical Medicine, Liverpool, UK

**Author notes:** These authors contributed equally (Marilene M. Ambadiang, Caroline Fouet).

**Keywords:** Anopheles, oviposition behavior, chemosensory receptor genes, ecological specialization, population genomics, urban adaptation

## Abstract

Oviposition site choice is a major determinant of habitat selection in insects, yet its behavioral and genetic bases remain poorly understood. Although model organisms such as *Drosophila melanogaster* provide well-established systems for studying oviposition, disentangling foraging from egg-laying decisions remains challenging because females use the same substrates for feeding and reproduction. Here, we identify a pair of sister mosquito species that exhibit contrasting oviposition strategies and provide a promising system for investigating how divergence in sensory information processing shapes behavioral shifts between generalist and specialist taxa. Using binary-choice assays, we tested oviposition preferences in 1,046 gravid females from both field and laboratory populations. *Anopheles gambiae* females were highly selective and deposited nearly all eggs in a single water source, consistent with specialist behavior. In contrast, females of its sibling species, *An. coluzzii*, adopted a generalist strategy marked by a more even distribution of eggs across available substrates. To identify chemosensory receptor genes that diverge between the cryptic species and may underlie these behavioral differences, we analyzed amino acid substitutions across odorant, ionotropic, and gustatory receptor families using whole-genome sequencing data. Although differentiation was limited at the gene-family level, several individual divergent loci associated with carboxylic acid, amine, and other volatile detection pathways emerged as candidate drivers of species-specific oviposition behavior. These findings suggest that shifts in sensory perception during egg laying may facilitate behavioral and ecological specialization, and that phenotypic divergence in environmental sensing can arise before substantial genome-wide differentiation.

## 1. Introduction

Behavioral divergence is a driver of adaptation and diversification, especially in species undergoing ecological segregation (Arnold, 1981; Heth & Nevo, 1981; Mayr, 1942, 1963; Pillay & Rymer, 2012). Over the past decades, studies on host shifts in insects have provided striking examples of how behavioral adaptations shape species evolution (Bush, 1969; Jaenike, 1990; Thorsteinson, 1960; Visser, 1986). Where insect females lay their eggs determines both offspring survival and habitat use. Consequently, oviposition site selection provides an important framework for understanding how behavioral variation promotes ecological specialization (Chess & Ringo, 1985; Jaenike, 1978; Richmond & Gerking, 1979).

Fundamental insights into oviposition decision-making have been gained from decades of research on saprophytic and herbivorous insects, in which generalist and specialist females often use the same resources for feeding and egg laying, creating overlapping cues that obscure the distinction between foraging and oviposition choices. (Amlou et al., 1998; Dekker et al., 2006; Jaenike, 1990; Joseph et al., 2009; Lihoreau et al., 2016; Liu et al., 2020; Schäpers et al., 2015; Thorsteinson, 1960; Vijayan et al., 2022; Visser, 1986; Weeraddana & Evenden, 2022; Wiklund, 1981). In mosquitoes, however, feeding and oviposition occur at different times and involve distinct resources, making them an attractive model for investigating the genetic basis and evolutionary underpinnings of oviposition behavior (Afify & Galizia, 2015; Bentley & Day, 1989; Day, 2016; Mwingira et al., 2020). Oviposition preference has been extensively studied in mosquitoes, particularly within the framework of the optimal oviposition theory, which posits that females select habitats that maximize the fitness of their offspring (Jaenike, 1978). Laboratory and semi-field experiments indicate that female mosquito oviposition decisions are determined by predator presence, conspecific density, competition, and abiotic factors (Allgood & Yee, 2018; Beehler et al., 1993; Blaustein et al., 2004; Blaustein & Kotler, 1993; Bond et al., 2005; Costa-da-Silva et al., 2024; Cozzer, 2019; Dhileepan, 1997; Lampman & R Novak, 2005; Obenauer et al., 2010; O’Meara et al., 1989). Despite this significant research, the extent to which oviposition behavior contributes to ecological specialization remains unexplored.

The *Anopheles gambiae* complex comprises a group of cryptic species with striking ecological and behavioral divergence, offering a suitable model for studying how behavioral shifts affect ecological adaptation at early stages of speciation (Coluzzi et al., 1977, 1979; Davidson & Jackson, 1962). The existence of freshwater- and saltwater-breeding variants of *An. gambiae* sensu lato was recognized as early as the 1940s (Muirhead-Thomson, 1946, 1951; Paterson, 1962). Incipient speciation was later confirmed in the 1960s when laboratory crosses between certain subpopulations produced sterile F₁ males, leading to the formal recognition of *An. gambiae* as a species complex (Davidson, 1964; Davidson & Jackson, 1962). The complex contains at least nine taxa including two saltwater-breeding species (*An. melas* and *An. merus*) and seven freshwater-breeding species (*An. gambiae* sensu stricto, *An. coluzzii*, *An. arabiensis*, *An. quadriannulatus*, *An. amharicus*, *An. bwambae*, and *An. fontenillei*) (Barrón et al., 2019; Coetzee et al., 2013; Riehle et al., 2011).

Within the species complex, *An. gambiae* and *An. coluzzii* are the most efficient vectors of malaria parasites. Despite extensive ongoing hybridization, the two morphologically indistinguishable species diverged approximately 0.35 million years ago and occupy distinct ecological niches at different geographic/spatial scales (Barrón et al., 2019; della Torre et al., 2002, 2005; Touré et al., 1998). In West and Central Africa, *An. gambiae* typically breeds in ephemeral, rain-dependent puddles in rural environments, while *An. coluzzii* is adapted to more permanent, human-made breeding sites in urban settings or in arid ecosystems (Costantini et al., 2009; Diabate et al., 2005; Edillo et al., 2002, 2006; Gimonneau et al., 2012; Kamdem et al., 2012; Simard et al., 2009). Even in sympatric zones, the abundance of immature stages differs within larval habitats likely due to variations in oviposition behavior and larval performance (i.e., survival, development, and growth in a given environment) between the two species (Edillo et al., 2002, 2006; Kamdem et al., 2012). Laboratory studies have revealed differences in larval performance between *An. gambiae* and *An. coluzzii*, including predator avoidance behaviors (Gimonneau et al., 2010; Roux et al., 2013, 2014), predator-mediated fitness (Niang et al., 2020), development time (Mouline et al., 2012), sensitivity to ammonium concentration (Tene Fossog et al., 2013), and susceptibility to agrochemicals (Ambadiang et al., 2024). Although habitat segregation has been well studied and linked to differential larval survival between the cryptic species, the role of oviposition site selection remains unknown.

To select suitable oviposition sites, female mosquitoes rely on neural and sensory-processing systems to interpret visual, olfactory, gustatory, and tactile signals from the environment (Afify & Galizia, 2015; Beehler et al., 1993; Konopka et al., 2021; Mwingira et al., 2020). In *Anopheles* mosquitoes, laboratory choice assays have demonstrated that oviposition preference can be affected by diverse factors, including salinity gradients (Foley & Bryan, 1999; Nwaefuna et al., 2019; Osborn et al., 2006), plant and microbial volatiles (Asmare et al., 2017; Lindh et al., 2008; Milugo et al., 2024; Wondwosen et al., 2016), water from *Anopheles*-preferred vs. *Culex*-preferred habitats (Sumba et al., 2004), habitat quality (Suh et al., 2016), manure-based fertilizers (Hardy et al., 2023), cow urine (Kweka et al., 2011), substrate color (Lowassari et al., 2023), larval density and stage, predators, and light conditions (Agyapong et al., 2014). These studies reveal the complexity of environmental and chemosensory signals that can shape oviposition decisions in *Anopheles* females. Mosquito olfaction has been relatively well studied due to its role in host seeking and pathogen transmission (Bowen, 1991; Carey et al., 2010; Konopka et al., 2021; G. Wang et al., 2010). However, evidence from *Aedes* and *Culex* mosquitoes suggests that contact chemostimulants and visual cues play roles as important as volatile odorants in oviposition behavior (Beehler et al., 1993; Trexler et al., 1998). For example, knockdown of the gustatory receptor Gr11 impairs the ability of *Aedes* females to locate and recognize oviposition sites, further supporting the involvement of taste-related genes in oviposition behavior (Zhao et al., 2024). These findings underscore the need for a comprehensive analysis that explores both olfactory and contact-based sensory pathways to better understand how mosquito infochemical perception modulates oviposition decisions.

In this study, we investigated the behavioral and genetic bases of divergent oviposition preferences in *An. gambiae* and *An. coluzzii*. Using binary-choice experiments, we characterized the degree of selectivity or flexibility in oviposition choices in both field-collected and laboratory-reared females. To explore the molecular basis of behavioral divergence between species, we analyzed genome-wide variation and amino acid substitutions across chemosensory gene families using pooled whole-genome sequencing (Pool-seq) and the *Anopheles gambiae* 1000 Genomes Phase 2 dataset (The Anopheles gambiae 1000 Genomes Consortium et al., 2020). Our results revealed contrasting oviposition behaviors in *An. gambiae* and *An. coluzzii* despite minimal detectable genome-wide differentiation. These findings suggest that behavioral specialization at early stages of ecological segregation in cryptic *Anopheles* species may be driven by sensory divergence acting on a small subset of chemosensory receptor genes.

## 2. Materials and Methods

### 2.1 Mosquito populations

Gravid females of *An. gambiae* and *An. coluzzii* were tested in two-choice oviposition experiments under laboratory conditions to assess preference between two types of water. We examined behaviors in field-collected mosquitoes and in two laboratory colonies: *An. coluzzii* Ngousso, established in 2006, and *An. gambiae* Kisumu, established in 1975. To assess oviposition preferences at early stages of ecological and genetic divergence, we tested sympatric populations from a small area around Yaoundé, Cameroon, and from surrounding rural environments (Fig.1). In this area, *An. coluzzii* occurs in densely urbanized areas and is absent from rural zones characterized by degraded forest habitats where *An. gambiae* predominates. Both species co-occur in suburban areas (Ashu et al., 2024; Kamdem et al., 2012, 2017; Tene Fossog et al., 2013). *Anopheles coluzzii* females were sampled from four urban neighborhoods of Yaoundé: Etoa Meki (3°52’48" N, 11°31’39" E), Combattant (3°53’01" N, 11°30’32" E), Ekié (3°49’58" N, 11°32’19" E), and Voirie Municipale (3°51’35" N, 11°31’11" E). *Anopheles gambiae* females originated from the suburban sites of Evogo (3°58’60" N, 11°35’60" E) and Nkolondom (3°56’43ʹʹ N, 11°30’01" E). Field populations were established from larvae collected in naturally occurring breeding sites. Larvae were reared to adults under standard insectary conditions (27 ± 1 °C, 75 ± 5% relative humidity, 12:12 h light:dark cycle), and adult females were blood-fed 3–4 days post-emergence and used for oviposition experiments.

**Figure 1:**
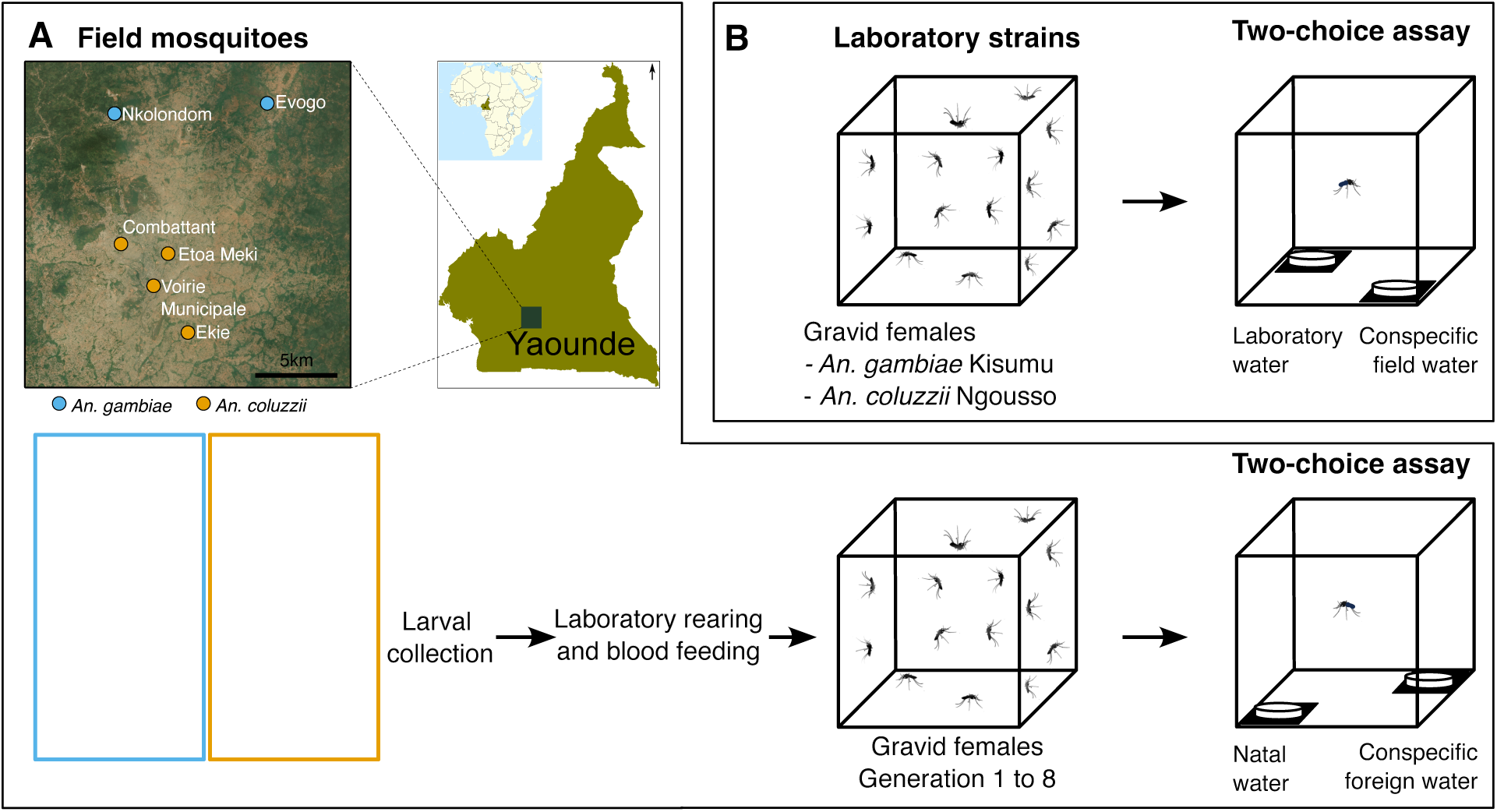
Experimental design for oviposition assays in field and laboratory populations. (A) Field populations. Gravid females originating from six sites around Yaoundé, Cameroon (Nkolondom, Evogo, Combattant, Etoa Meki, Voirie Municipale, and Ekié) were tested in two-choice oviposition assays. The map shows sampling locations for *Anopheles gambiae* and *An. coluzzii*, and photographs illustrate typical larval habitats from which larvae and natal water were collected. Field-collected larvae were reared in the laboratory to obtain gravid females (generations F1–F8), which were then exposed to dual-choice assays comparing natal water with conspecific foreign water. (B) Laboratory strains. Gravid females from the laboratory colonies *An. gambiae* Kisumu and *An. coluzzii* Ngousso were tested using the same two-choice assay design, with oviposition choices between laboratory water and conspecific field water.

### 2.2 Two-choice oviposition experiment

Gravid females (72–96 h post-blood meal), including those from the two laboratory colonies (Ngousso and Kisumu) and wild populations (F1–F8 generations), were individually tested for oviposition behavior (Fig.1). Egg counts in choice experiments provide a simple yet robust approach widely used to test oviposition behavior in insects (Álvarez-ocaña et al., 2023; Chess & Ringo, 1985; Day, 2016; Okal et al., 2015). Two-choice tests aim to assess the selectivity of females when presented with two distinct oviposition substrates. We used two-choice experiments to quantify the ability of *Anopheles* mosquitoes to consistently discriminate between waters from different origins. For field-collected mosquitoes, each female was offered a choice between natal water and conspecific foreign water. Natal water was collected from the original breeding site where larvae were sampled, or from an adjacent site within 30 meters. Conspecific foreign water was obtained from a site located at least 3 km away, within the known distribution of the species (urban areas for *An. coluzzii* and rural or suburban areas for *An. gambiae*, Fig. 1, Supplementary Materials Table S1).

Laboratory strains were offered a choice between laboratory-rearing water and natural water from breeding habitats of conspecific wild populations (Fig.1). All assays using field-collected females were conducted from June 2021 to June 2023 during the rainy seasons when natural breeding sites were abundant. Water was collected on the day of the experiment using dippers from confirmed *Anopheles*-positive sites, filtered through 0.5 μm mesh to remove debris, and used immediately. Each gravid female was placed overnight in a 30 × 30 × 30 cm mesh cage under insectary conditions. To minimize odor contaminations between replicates, the cage and the mesh were washed between experiments. Two identical Petri dishes (5 cm diameter × 1 cm height) were positioned equidistant from the cage center. One contained natal or laboratory water and the other was filled with an equal volume (3 ml) of conspecific foreign or field water. To minimize positional bias, the two petri dishes were randomly placed in the cage, diagonally on black square paper (10 × 10 cm) to enhance visual contrast (Fig. 1). A 10% sugar solution was provided via cotton pads placed on top of the cage. Eggs were counted using a wireless digital microscope the following day after a 24-hour oviposition period.

### 2.3 Oviposition metrics

We calculated four behavioral metrics from individual egg counts to quantify oviposition performance and selectivity. These metrics were compared across the four populations in which at least 100 females laid eggs in the choice assays: *An. gambiae* field, *An. coluzzii* field, *An. gambiae* laboratory (Kisumu), and *An. coluzzii* laboratory (Ngousso). Field-collected females from multiple sites were pooled within each species to ensure adequate sample size per population. Oviposition rate was defined as the proportion of females that laid eggs out of the total number tested, and fecundity was measured as the total number of eggs laid per female. Oviposition behavior was further characterized using two additional indices.

First a split ratio was calculated for each female as the number of eggs laid in the less-preferred water divided by the total number of eggs laid in both petri dishes. This ratio measures the evenness of egg distribution between the two oviposition substrates. A value of 0 indicates exclusive oviposition in one petri dish, while 0.5 indicates an equal split. Oviposition preference was then quantified using the Oviposition Preference Index (OPI), defined as OPI = (Nn − Nf)/(Nn + Nf), where Nn is the number of eggs laid in natal (or laboratory) water and Nf is the number laid in foreign (or conspecific field) water. This index was calculated only for females exhibiting a strong preference (i.e., ≥70% of eggs deposited in a single petri dish). This approach was used to ensure that the resulting index reflects a deliberate choice or aversion, filtering out individuals who may be indifferent or whose egg-laying behavior is less consistent (Dawkins, 1969; Hopkins et al., 1999; Monks & Kelly, 2003; Withers, 1999). Positive OPI values indicate preference for natal (or laboratory) water, while negative values indicate preference for foreign (or conspecific field) water. All analyses were conducted in R v4.3.1 (R Core Team, 2016).

### 2.4 Genome-wide differentiation between species

To assess the extent of genomic differentiation among populations that were tested in oviposition preference assays, we performed pooled whole-genome sequencing (Pool-seq) on field-collected samples. Wild larvae were collected from breeding sites in the five sampling locations (Nkolondom, Evogo, Ekié, Voirie Municipale, and Combattant) and reared to adults in the insectary. Adult females were identified to species using PCR–RFLP assays performed on DNA extracted from legs (Fanello et al., 2002). Individuals identified as *An. gambiae* or *An. coluzzii* were then pooled for sequencing. For each population, total genomic DNA was extracted from 2–5-day-old female mosquitoes using the DNeasy Blood & Tissue Kit (Qiagen, Germany), following the manufacturer’s protocol. DNA quality was assessed by gel electrophoresis and quantified using a Qubit 4 Fluorometer (Thermo Fisher Scientific). Equal amounts of DNA from each individual were pooled to generate one library per population. Each pool consisted of 100 mosquitoes, contributing approximately 10 ng of DNA per individual. Sequencing libraries were constructed using the NEBNext Ultra II DNA Library Prep Kit (New England Biolabs). Paired-end sequencing (150 bp reads) was performed on an Illumina NovaSeq 6000 platform (Novogene) to a target depth of ∼80× per population.

Raw reads were quality-checked with FastQC, trimmed using Trimmomatic (Bolger et al., 2014), and aligned to the *Anopheles gambiae* PEST reference genome (AgamP4) using BWA v0.7.13 (Li & Durbin, 2009). PCR duplicates were marked using Picard v2.1.1, and uniquely mapping reads with high mapping quality were retained for downstream analyses. Variants were called using the *mpileup* function in Samtools v1.3 (Li et al., 2009), and SNPs were synchronized and filtered with PoPoolation2 v1.201 (Kofler et al., 2011) using default parameters: minimum base quality of 20, minimum coverage of 20, and a minor allele count threshold of 2. The resulting .sync file was used to estimate *F*_ST_ and conduct genome-wide scans of population differentiation.

We estimated pairwise *F*_ST_ using two complementary approaches implemented in PoPoolation2 and poolfstat v2.0 (Hivert et al., 2018) respectively. For poolfstat, *F*_ST_ was computed from allele count data using an ANOVA-based estimator. PoPoolation2 *F*_ST_ estimates were generated using a sliding window approach. *F*_ST_ values from both methods were smoothed using Gaussian kernel smoothing (bandwidth = 300 kb) with the *locpoly()* function from the *KernSmooth* R package.

### 2.5 Diversity and differentiation across chemosensory receptor genes

To examine genetic diversity and differentiation among receptor gene families involved in chemosensory perception, we analyzed nonsynonymous variation in odorant (ORs), ionotropic (IRs), and gustatory receptors (GRs). These gene families form ligand-gated ion channels that mediate behaviors such as host-seeking, oviposition site selection, and blood-feeding (Carey et al., 2010; Chen & Amrein, 2017; McBride et al., 2007; Raji & Potter, 2022; Sánchez-Gracia et al., 2009; Vosshall et al., 1999). *Anopheles* mosquitoes possess ∼75 OR, ∼40 GR and ∼80-90 IR genes (Giraldo-Calderon et al., 2015). In this study, we selected 23 OR genes based on prior evidence of strong excitatory responses to chemical stimuli, including known oviposition attractants (Carey et al., 2010). In parallel, we screened 39 IR genes based on functional annotation in *An. gambiae* including candidates responsive to amines and carboxylic acids (Pitts et al., 2017). For GRs, we analyzed 29 genes identified via phylogenetic classification and transcriptomic profiling of appendage-specific tissues in *An. coluzzii*, *An. gambiae*, and *Aedes aegypti* (Carr et al., 2021).

We assessed genetic variation using the *Anopheles gambiae* 1000 Genomes Phase 2-AR1 dataset, which includes whole-genome sequences from 1,142 wild-caught mosquitoes sampled from 33 sites in 13 sub-Saharan African countries (Miles et al., 2017; The Anopheles gambiae 1000 Genomes Consortium et al., 2020). Sequencing was performed using 100-bp Illumina reads at an average depth of ∼30× per individual. Read alignment was carried out using BWA v0.6.2 (Li & Durbin, 2009), and variant calling was performed with the GATK UnifiedGenotyper (v2.7.4) (McKenna et al., 2010). For our analysis, we selected a subset of 532 individuals identified as *An. gambiae* (n = 332) or *An. coluzzii* (n = 189) from 10 countries representing the geographic diversity of sub-Saharan Africa (Table S2).

We retrieved exon coordinates for the selected chemosensory genes from the *An. gambiae* Agam P4 genome annotation (GFF) file using accession numbers (Giraldo-Calderon et al., 2015). Using bcftools v1.21 (Li et al., 2009), we parsed VCF files of the selected 532 individuals to extract variants mapping to exons of each gene retaining only those with minor allele frequency (MAF) > 0.01. Nonsynonymous mutations were annotated and extracted from VCF files using two different programs: SnpEff (Cingolani, Patel, et al., 2012; Cingolani, Platts, et al., 2012) with the prebuilt *An. gambiae* database and the Ensembl Variant Effect Predictor (VEP v115) (McLaren et al., 2016). To analyze diversity and divergence among the selected genes, we included only variants classified as missense and biallelic by both annotation programs.

To assess genetic diversity, we calculated the observed heterozygosity or the frequency of heterozygous genotypes for each variant within each gene in *An. gambiae* and *An. coluzzii*. We then estimated the mutation load per gene by deriving the proportion of variant sites bearing heterozygous genotypes but no homozygous mutant allele (heterozygous exclusive) as well as the frequency of sites with homozygous mutant alleles. To evaluate differentiation between *An. gambiae* and *An. coluzzii*, we calculated *F*_ST_ per variant site using Hudson’s estimator in PLINK 2.0 (Chang et al., 2015; Hudson et al., 1992).

## 3. Results

### 3.1 Oviposition rate and fecundity vary between species and long-term colonies

A total of 1,046 gravid *Anopheles* females were exposed to two-choice oviposition experiments, with 20–40 individuals tested per replicate. Each replicate included gravid females released individually in cages containing the two types of water. Among the 1,046 females tested, 438 laid eggs within 24 hours. Oviposition rates were comparable between field populations of *An. coluzzii* (110/327 = 33.6%) and *An. gambiae* (123/370 = 33.2%). In contrast, a substantially higher percentage of females from laboratory strains laid eggs compared to their conspecific field populations: *An. coluzzii* Ngousso (100/165 = 60.7%) and *An. gambiae* Kisumu (105/184 = 57.1%) (χ² = 0.316, df = 1, p = 0.574) (Fig. 2, Table S1).

**Figure 2.**
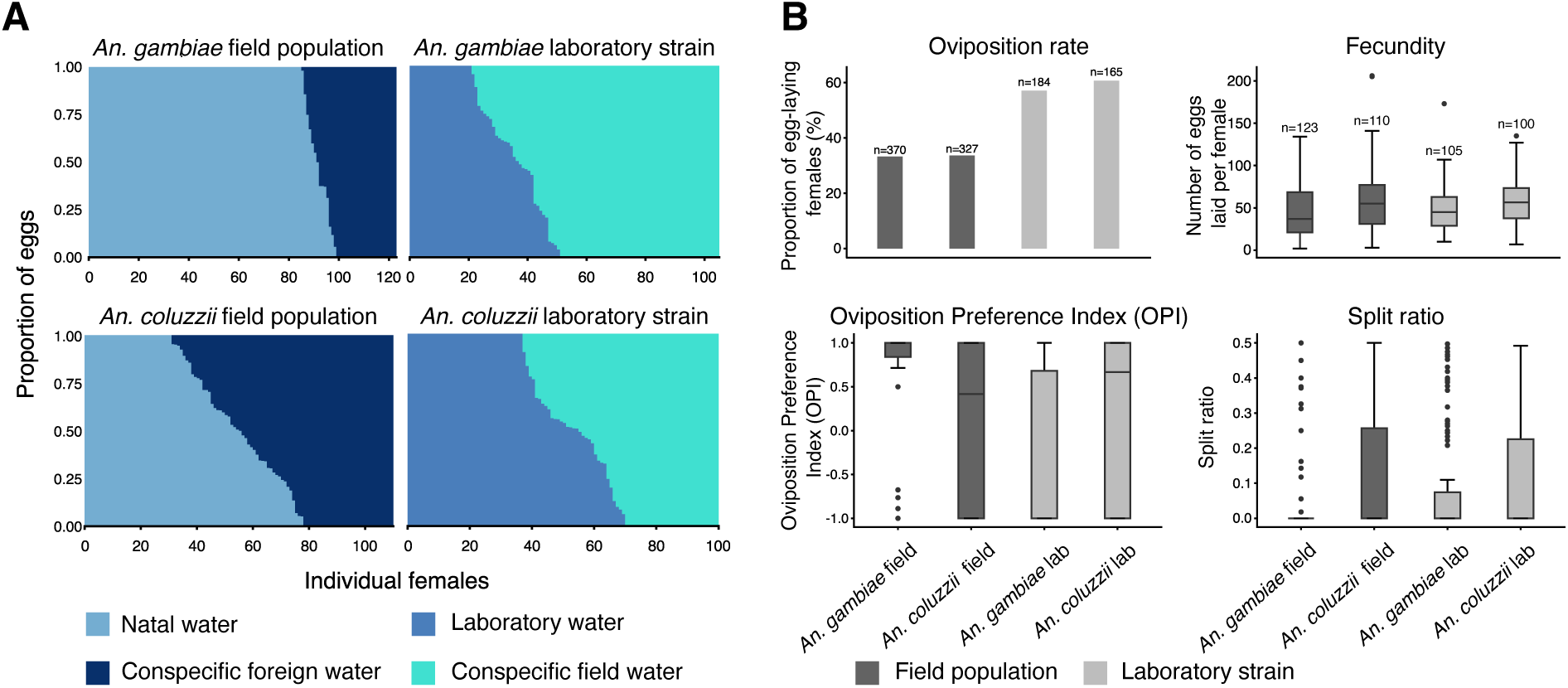
Contrasting oviposition behaviors in *Anopheles gambiae* and *An. coluzzii* from field populations and laboratory colonies. (A) Egg-laying patterns. Distribution of eggs laid by individual gravid females tested in dual-choice assays. Field-collected females were exposed to natal water and conspecific foreign water, whereas laboratory colonies were tested with laboratory water versus conspecific field water. Colors correspond to the four water types displayed in the legend. (B) Oviposition metrics across populations. Barplots and boxplots summarize oviposition rate (proportion of females laying eggs), fecundity (number of eggs per female), oviposition preference index (OPI), and split ratio. Laboratory colonies of both species exhibited higher oviposition rates than field populations (χ² = 0.316, df = 1, *p* = 0.574). Pairwise Wilcoxon tests show that the *An. coluzzii* laboratory strain laid significantly more eggs than all other populations (*p* < 0.05). Field-collected *An. gambiae* populations showed very restrictive oviposition behavior, with minimal egg splitting and OPI values close to +1, indicating a strong preference for natal water.

Among females that laid eggs (n = 438), we compared the number of eggs laid per individual (fecundity), focusing on the four main populations for which at least 100 females laid eggs (*An. gambiae* field, *An. gambiae* lab, *An. coluzzii* field and *An. coluzzii* lab). Fecundity differed significantly among populations (Kruskal–Wallis χ² = 16.30, df = 3, p = 9.84 × 10⁻⁴; Fig. 2). Mean fecundity (± 95% CI) was 58.4 eggs/female/24 hours (51.8–65.0) for *An. coluzzii* field, 58.7 (53.2–64.2) for *An. coluzzii* lab, 46.8 (41.2–52.5) for *An. gambiae* field, and 48.7 (43.8–53.6) for *An. gambiae* lab. Pairwise Wilcoxon comparisons indicated that both *An. coluzzii* populations laid significantly more eggs than the *An. gambiae* field population (p = 0.0115 and p = 0.0043, respectively). No significant differences were detected between the two *An. coluzzii* populations (p = 0.608) or between the two *An. gambiae* populations (p = 0.268). These results indicate that fecundity under the standardized two-choice conditions is shaped by both species identity and environmental origin, with *An. coluzzii*, specifically the laboratory strain, laying substantially more eggs than *An. gambiae*.

### 3.2 *Anopheles coluzzii* distributes eggs more evenly among water sources than *An. Gambiae*

To quantify flexibility in egg distribution among females, we calculated a split ratio for each individual, defined as the proportion of eggs deposited in the less-preferred water. Split ratios ranged from 0 to 0.5 among tested females (Fig. 2). As with fecundity, the split ratio differed significantly between the four populations (Kruskal–Wallis: χ² = 28.65, df = 3, *p* = 2.6 × 10⁻⁶). Mean ratios (± 95% CI) were 0.120 (0.090–0.153) for *An. coluzzii* field, 0.112 (0.078–0.148) for *An. coluzzii* lab, 0.031 (0.015–0.050) for *An. gambiae* field, and 0.084 (0.055–0.114) for *An. gambiae* lab. Pairwise Wilcoxon tests indicated that both *An. coluzzii* populations had significantly higher split ratios than the *An. gambiae* field population (*p* = 5.3 × 10⁻⁷ and 1.4 × 10⁻⁴, respectively). *Anopheles gambiae* lab also split eggs more frequently than *An. gambiae* field (*p* = 0.002). No difference was detected between the two *An. coluzzii* populations (*p* = 0.374). The *An. gambiae* laboratory colony (Kisumu) was also very selective compared to *An. coluzzii* populations but showed a relatively higher level of flexibility relative to conspecific field populations. When fecundity was considered, split ratio was weakly but significantly positively correlated with the total number of eggs laid (Spearman ρ = 0.234, *p* = 7.6 × 10⁻⁷). This suggests that females with higher fecundity were slightly more likely to distribute eggs across both oviposition sites. These findings indicate that *An. gambiae* field females prefer a strong commitment to a single oviposition site, whereas *An. coluzzii* populations have a greater tendency to distribute eggs across the available choices. Overall, species identity emerged as the primary determinant of oviposition flexibility in the cryptic species. *Anopheles coluzzii* was characterized by high split ratios, consistent with a bet-hedging strategy, whereas *An. gambiae*, particularly the field population, tended to deposit eggs in a single site, suggesting a more selective oviposition behavior.

### 3.3 The cryptic species exhibit specialist and generalist oviposition behaviors

In addition to split ratios, we assessed oviposition choices using the Oviposition Preference Index (OPI), restricting analyses to females showing strong preference (≥70% of eggs deposited in a single dish). The OPI ranges from −1 (complete preference for the foreign water or conspecific field water) to +1 (complete preference for the natal water or laboratory water), with 0 indicating no preference. We found significant differences in OPI values among the four populations (Kruskal–Wallis χ² = 50.9, df = 3, *p* = 5.1 × 10⁻¹¹, Fig.2). Pairwise Wilcoxon tests showed that both *An. gambiae* populations have highly polarized oviposition choices in the experimental conditions (*p* ≤ 0.001 for all comparisons involving *An. gambiae* field vs the two *An. coluzzii* populations; *p* = 3.5 × 10⁻¹¹ for *An. gambiae* field vs *An. gambiae* lab). Field-collected *An. gambiae* females, including individuals reared in laboratory conditions for 1–8 generations, overwhelmingly preferred their natal field water (median OPI = 1). *Anopheles gambiae* laboratory females (Kisumu) also strongly preferred conspecific field water (median OPI = −1), indicating that even after long-term colonization, this strain has retained the ability to detect and select water sources characteristic of natural breeding sites.

In contrast, both field-collected and laboratory-reared *An. coluzzii* females showed broad OPI values, with no significant difference between populations (pairwise Wilcoxon: *p* = 0.51), consistent with split-ratio results indicating flexible oviposition preferences. In summary, fecundity, egg-splitting behavior, and oviposition preference index identify *An. gambiae* as an oviposition specialist characterized by relatively lower fecundity and more restrictive breeding site selection whereas *An. coluzzii* exhibits a generalist strategy with higher fecundity and more flexible oviposition choices.

### 3.4 Genome-wide differentiation is low both within and between species

Using Pool-seq, we assessed genetic differentiation across five *Anopheles* populations from Yaoundé and surrounding areas (Nkolondom, Ekié, Evogo, Combattant, and Voirie Municipale), all of which were included in oviposition choice assays. A total of 11,543 SNPs shared among the five populations were used to compute pairwise *F*_ST_ after filtering for coverage (20–500 reads per SNP per pool) and a minor allele frequency (MAF) ≥ 0.001. Genome-wide *F*_ST_ values were globally low, indicating reduced genetic structure at this geographic scale (*F*_ST_ < 0.12). Among *An. coluzzii* populations, pairwise *F*_ST_ ranged from 0.002 to 0.006, while *An. gambiae* populations showed a mean *F*_ST_ of 0.004 ± 0.002 (Table 1). Comparisons between *An. coluzzii* and *An. gambiae* revealed *F*_ST_ values more than ten-fold higher relative to those observed within species, with mean interspecific *F*_ST_ ranging from 0.107 ± 0.006 to 0.116 ± 0.006. These values indicate a detectable level of genetic isolation between the cryptic species in the sympatric region of Yaoundé.

**Table 1:**
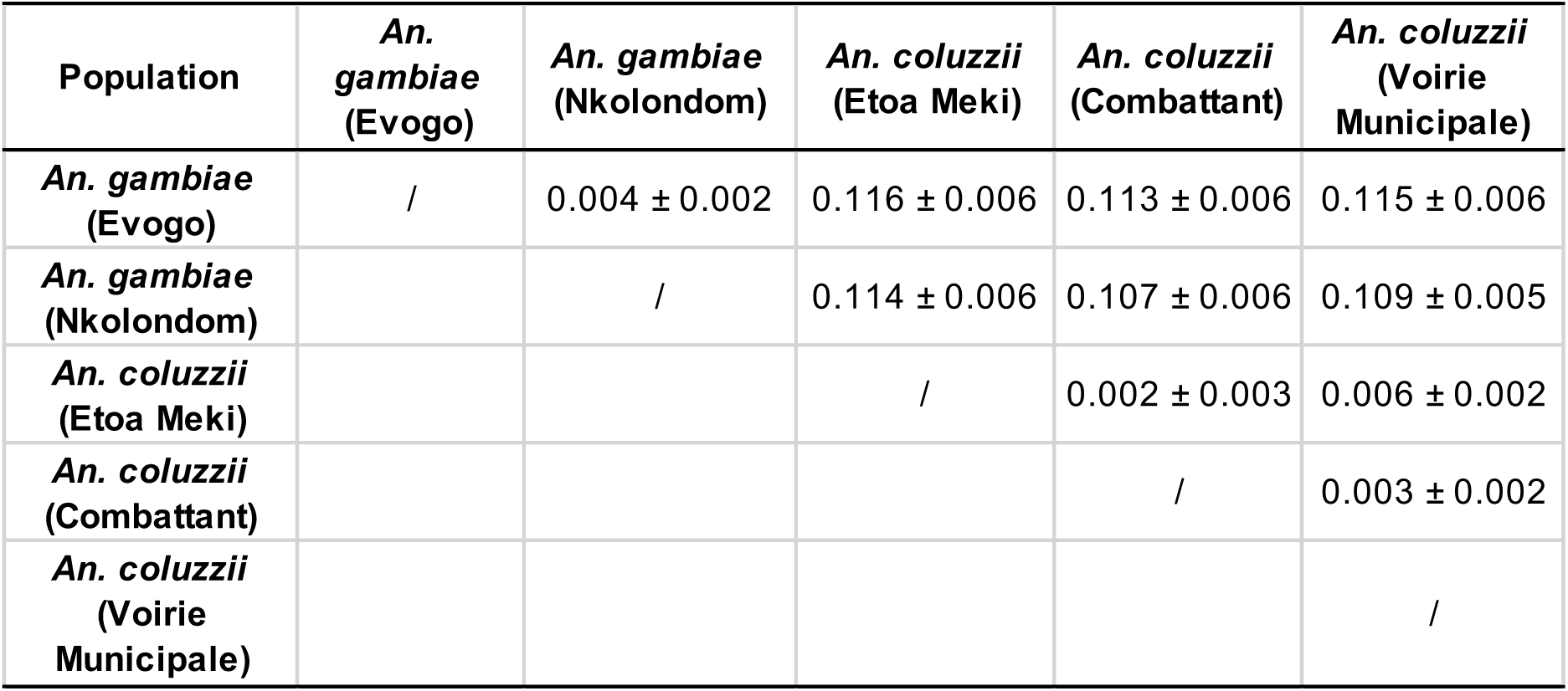
Genome-wide *F*_ST_ estimated with poolfstat. Standard errors were obtained using block-jackknife resampling.

To identify genomic regions of high differentiation, we performed genome scans of *F*_ST_ values using 300-kb sliding windows across the five chromosome arms (Fig. 3). Outliers were defined as windows exceeding the 99.9^th^ percentile of the genome-wide *F*_ST_ distribution. Both poolfstat and PoPoolations2 revealed consistent patterns of *F*_ST_ across the genome (Fig. 3). Therefore, outlier detection was conducted using PoPoolations2-derived *F*_ST_ estimates only.

**Figure 3.**
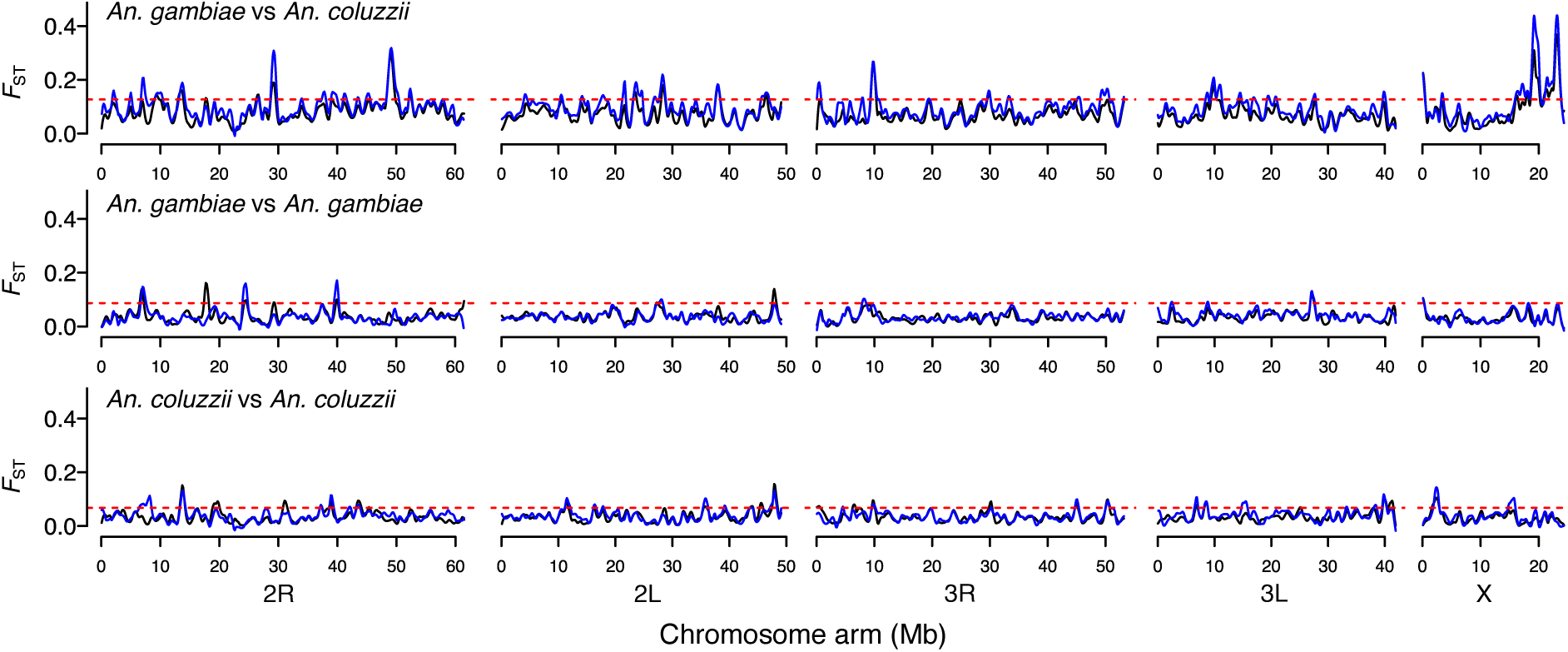
Genome-wide patterns of genetic differentiation across the five chromosome arms. Pairwise *F*_ST_ values were calculated in 300-kb sliding windows for three comparisons: (top) *An. gambiae* Nkolondom vs *An. coluzzii* Voirie Municipale, (middle) *An. gambiae* Nkolondom vs *An. gambiae* Evogo, and (bottom) *An. coluzzii* Combattant vs *An. coluzzii* Voirie Municipale. For each comparison, *F*_ST_ estimates from poolfstat (black line) are overlaid onto values obtained with PoPoolation2 (blue line) to highlight concordant regions of significant differentiation. The horizontal dashed line denotes the 99.9^th^ percentile threshold used to identify genomic outlier windows.

Among the three *An. coluzzii* populations, we performed pairwise comparisons and retained only SNPs that appeared as outliers in all comparisons. This conservative approach identified 15 to 40 outlier SNPs, but none was shared between all three comparisons (Table S3). We then compared the two *An. gambiae* populations (Nkolondom and Evogo) and identified five outlier SNPs including one variant that mapped to AGAP029236, a gene predicted to encode metabotropic glutamate receptor 1, which plays a role in neural signaling (Fig. 3). The other four SNPs occurred in outlier windows also found in *An. coluzzii* comparisons (Table S3).

Between *An. gambiae* and *An. coluzzii*, we performed six pairwise comparisons and identified 8 to 50 outlier SNPs in each case (Fig. 3, Table S3). Among these, three SNPs were detected in all comparisons, including two variants within AGAP001084, encoding a fatty acyl-CoA reductase, and one SNP in AGAP001082, annotated as a saposin domain-containing protein (Table S3).

### 3.5 Mutation load in chemosensory gene families is higher in *An. gambiae* than in *An. Coluzzii*

To investigate genetic diversity within chemosensory gene families and among species, we estimated observed heterozygosity as the frequency of heterozygous genotypes for each mutation. To compare the mutation loads, we computed the proportion of exclusively heterozygous mutations (i.e., the fraction of variants exhibiting heterozygous genotypes but no homozygous mutant individuals) and the fraction of variants bearing homozygous mutant (alternate) alleles. All three statistics were derived for each gene within the IR, GR, and OR families using biallelic variants jointly annotated as missense by SnpEff and Ensembl VEP from the *Anopheles gambiae* 1000 Genomes Project dataset (*An. gambiae*, n = 332 and *An. coluzzii*, n = 189). Nonsynonymous variation was widespread in chemosensory receptor gene families (Fig. 4). The IR family harbored the largest overall number of biallelic missense variants (1,870 on 39 genes), with a median of 47 sites per gene (Interquartile range, IQR = 31–59; range = 8–98), indicating a relatively even mutational burden per gene. GRs comprised 29 genes with 947 biallelic nonsynonymous mutations but a more skewed distribution (median = 33; IQR = 20-44; range = 5-71), including several genes with a very high number of mutations such as AgGr30 (71 missense variants). ORs included the fewest genes (23) and the smallest total of missense variants (675), with the lowest mutation load (median = 26; IQR = 18-43; range = 5–75), suggesting comparatively stronger evolutionary constraint in this family. The number of missense variants per gene differed significantly among the three families (Kruskal–Wallis χ² = 12.59, df = 2, *p* = 0.0018). Post-hoc Wilcoxon tests confirmed that IR genes contained significantly more missense variants per gene than both GRs (p = 0.010) and ORs (p = 0.005), whereas GRs and ORs did not differ (p = 0.461).

**Figure 4:**
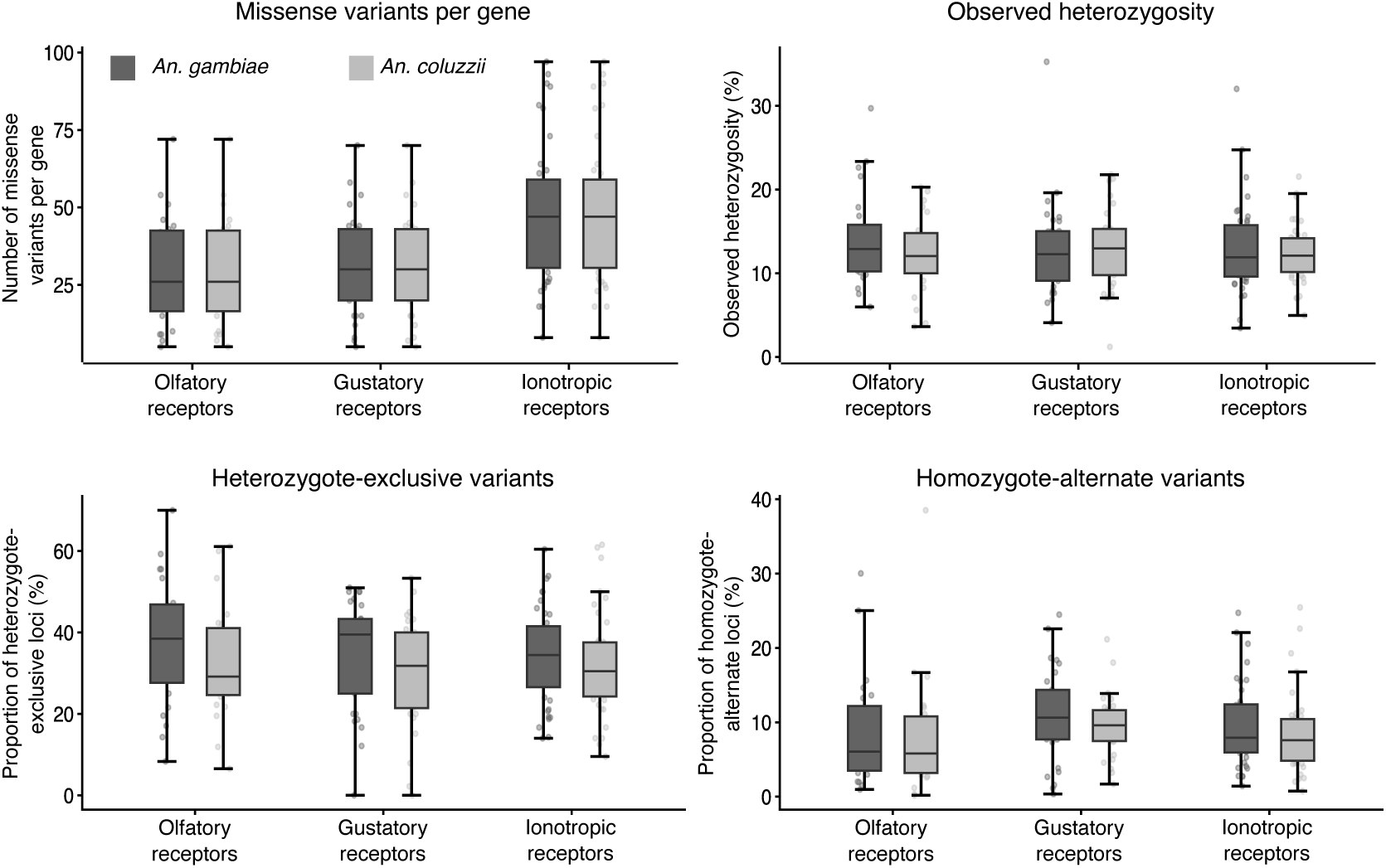
Mutation load and genetic diversity of missense variants across chemosensory receptor gene families. Boxplots summarize the number of biallelic missense variants per gene and the distribution of diversity metrics for ionotropic (IR), gustatory (GR), and odorant (OR) receptors in *An. gambiae* and *An. coluzzii*. The median number of missense variants per gene was slightly higher in IR genes compared with GR and OR, whereas observed heterozygosity did not differ significantly among gene families (Kruskal–Wallis test, *p* > 0.05). Both heterozygote-exclusive variants and homozygous alternate-allele variants occurred at higher proportions in *An. gambiae* than in *An. coluzzii* (paired Wilcoxon tests, *p* < 0.05), indicating a greater missense mutation load in *An. gambiae* across chemosensory receptor genes.

Observed heterozygosity (HZ) was similar among the three chemosensory gene families within each species, with no significant differences detected in either *An. gambiae* (Kruskal–Wallis *p* = 0.88) or *An. coluzzii* (*p* = 0.62, Fig. 4). In cross-species comparisons, HZ did not differ between *gambiae* and *coluzzii* for GR or OR variants (paired Wilcoxon *p* = 0.52 and *p* = 0.058, respectively), whereas IRs showed a small but significant increase in heterozygosity in *An. coluzzii* (p = 4.3 × 10⁻⁴). The proportion of heterozygote-exclusive sites did not differ among families in either species as well (Kruskal–Wallis tests, *p* > 0.2 for all comparisons, Fig. 4). However, *An. gambiae* had slightly higher frequencies of heterozygote-exclusive mutations than *An. coluzzii* for all three families: paired Wilcoxon tests for IR (*p* = 0.0407), GR (*p* = 0.0161), and OR (*p* = 0.0122). Likewise, the proportion of sites bearing homozygous alternate genotypes was significantly higher in *An. gambiae* than in *An. coluzzii* for the IR and GR families (paired Wilcoxon tests: IR, *p* = 0.0011; GR, *p* = 0.022, Fig. 4), whereas no difference was detected for OR genes (*p* = 0.52). Thus, *An. gambiae* harbors a higher mutation load of missense polymorphism in both heterozygous and homozygous states. This finding suggests that differences in effective population size, demographic history, or selective constraints shape distinct mutation burdens across chemosensory receptor gene families between the two species (Miles et al., 2017).

### 3.6 Candidate genes of chemosensory differentiation within families

To characterize genetic differentiation in chemosensory receptor genes, we analyzed *F*_ST_ between *An. gambiae* and *An. coluzzii* both within and among the three receptor families. Using the same dataset of biallelic missense variants applied in the genetic diversity analyses, we estimated per-SNP *F*_ST_ for each of the 91 chemosensory receptor genes. Mean *F*_ST_ values were low (< 0.18) across genes, consistent with the weak genome-wide differentiation revealed between *An. gambiae* and *An. coluzzii* using Pool-seq (Fig. 5). *F*_ST_ values did not differ significantly among the IR, GR, and OR families (Kruskal–Wallis χ² = 2.79, df = 2, *p* = 0.25). Average *F*_ST_ values per gene ranged from 0.054 ± 0.005 (IRs) to 0.066 ± 0.006 (GRs), with broadly overlapping interquartile ranges. Pairwise Wilcoxon tests confirmed the absence of significant differences between GRs and IRs (*p* = 0.31), GRs and ORs (*p* = 0.73), or ORs and IRs (*p* = 0.46, Fig. 5).

**Figure 5.**
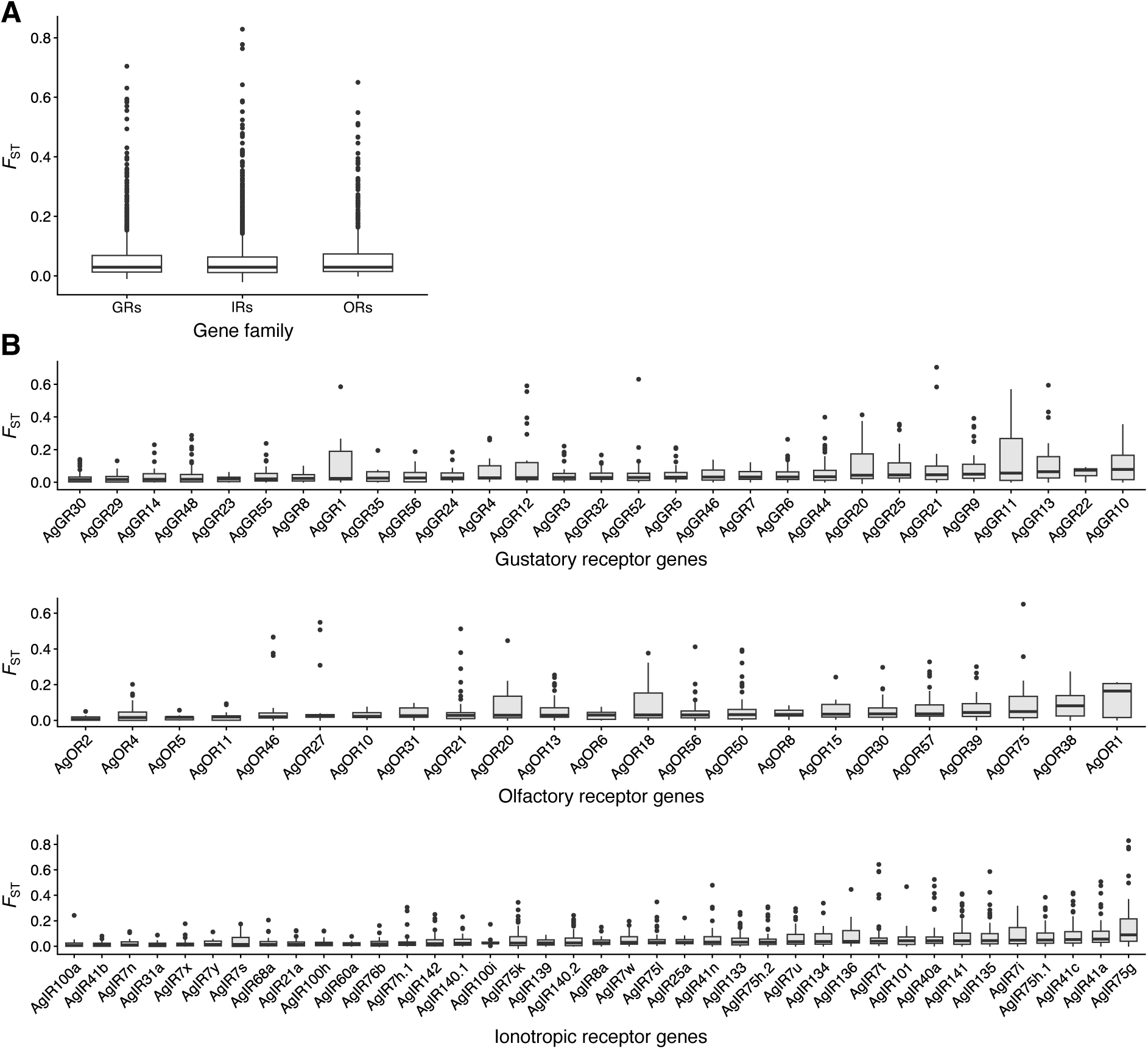
Genetic differentiation of chemosensory receptor genes between *Anopheles gambiae* and *An. coluzzii*. (A) Family-level differentiation. *F*_ST_ estimated from missense variants shows no significant difference in overall differentiation among ionotropic, gustatory, and odorant Receptors (Kruskal–Wallis χ² = 2.79, df = 2, *p* = 0.25). (B) Gene-level variation. Boxplots illustrate the distribution of per-SNP *F*_ST_ estimates, with median values per gene ordered from lowest to highest within each family. Despite limited divergence at the gene-family level, several individual genes show substantial median *F*_ST_ values and an accumulation of highly differentiated missense variants, suggesting they may contribute significantly to sensory divergence between the two species.

Nevertheless, the distribution of median *F*_ST_ values per gene was not uniform within families and showed a relatively skewed pattern in which a few loci exhibited substantial median *F*_ST_ and/or contained several variants with very high *F*_ST_ (Fig. 5). To identify the loci contributing most to differentiation within each family, we ranked the 91 chemosensory genes by their median *F*_ST_ values. The highest median *F*_ST_ highlighted five candidate loci as focal points of differentiation that may contribute to divergence in behaviors such as oviposition preference or host-seeking between *An. gambiae* and *An. coluzzii*: AgOr1 (median = 0.164, n = 5 variants), AgIr75g (0.089, n = 52), AgOr38 (0.082, n = 41), AgGr10 (0.078, n = 44), and AgGr22 (0.074, n = 5). Within each family, the four most differentiated genes based on median *F*_ST_ values were AgGr10, AgGr22, AgGr13, and AgGr11 among GRs; AgIr75g, AgIr41a, AgIr41c, and AgIr75h.1 among IRs; and AgOr1, AgOr38, AgOr75, and AgOr39 among ORs. The fourth gene with the highest median *F*_ST_ within the GR family, AgGr11, is a gustatory receptor gene implicated in the detection of water-borne cues during oviposition in *Aedes* mosquitoes (Zhao et al., 2024). Within the IR family, the known co-receptors (AgIr8a, AgIr25a, and AgIr76b) fell in the lowest range of differentiation, consistent with the idea that they play an essential role in physiological responses of ionotropic receptors and are therefore more constrained among species (Chen & Amrein, 2017; Sánchez-Alcañiz et al., 2018). Unsurprisingly, the most differentiated IRs were all tuning receptors, including AgIr41a and AgIr75g, which may be diverging in response to the diversity of ligands in each species’ ecological niche.

As a complementary approach to median *F*_ST_ per gene, we used the 99^th^ percentile of *F*_ST_ (*F*_ST_ ≥ 0.432) to detect outlier variants contributing to differentiation between *An. gambiae* and *An. coluzzii*. This analysis revealed nineteen genes harboring at least one outlier SNP across the three chemosensory receptor families. The strongest signal was found at AgIR75g, which contained six outlier SNPs, reaching *F*_ST_ values up to 0.829. Several additional IR loci had multiple outliers, including AgIR7t (3 sites, maximum *F*_ST_ = 0.642), AgIR40a (3 sites, maximum *F*_ST_ = 0.524), AgIR41a (2 sites, maximum *F*_ST_ = 0.507), and AgIR41n, AgIR101, and AgIR136 (one outlier each). Within the GR family, AgGR11 contained three outlier variants (maximum *F*_ST_ = 0.570), while AgGR21, AgGR12, AgGR52, AgGR13, and AgGR1 each carried one to two outliers. Among ORs, five loci (AgOR27, AgOR46, AgOR21, AgOR75, and AgOR20) harbored 1-2 outlier SNPs each (maximum *F*_ST_ = 0.650). By simultaneously considering median *F*_ST_ and the number of outlier variants, differentiation analyses point to AgIR75g as the most divergent locus, with AgIR41a, and AgGR11 forming a second tier of genes likely contributing to sensory shifts between *An. gambiae* and *An. coluzzii*.

## 4. Discussion

### 4.1 Oviposition preference reveals specialization in *An. gambiae*

In this study, we described differences in oviposition behavior between the cryptic species *An. coluzzii* and *An. gambiae*. *Anopheles coluzzii* females laid significantly more eggs than *An. gambiae* females under laboratory conditions. Based on research conducted in herbivorous insects, it has been suggested that behavioral specialists generally have lower fecundity compared to generalists (Mitter et al., 1993). The evolutionary explanation of the reduced fecundity of specialists remains unclear and could be linked to variables such as larger egg size, greater tendency to retain eggs, or non-adaptive factors (e.g., pleiotropic effects of mutations in genes underlying other adaptations) (Álvarez-ocaña et al., 2023; R’kha et al., 1997). The link between behavioral specialization and fecundity has been well-studied in *Drosophila* species where dietary generalists like *D. melanogaster* and *D. simulans* exhibit higher egg output than *D. sechellia*, which is an obligate specialist that feeds and reproduces almost exclusively on the ripe fruit of the noni plant (*Morinda citrifolia*) (Álvarez-ocaña et al., 2023; Auer et al., 2020; R’kha et al., 1997). In the present study, the difference in fecundity between the sibling species *An. gambiae* and *An. coluzzii* may reflect generalist/specialist contrasts, but complementary analyses such as egg retention and egg size distribution are still needed to support this hypothesis.

Skip-oviposition behavior differed significantly between *An. gambiae* and *An. coluzzii*, further highlighting the contrast in their reproductive strategies. Both field-collected and colonized *An. coluzzii* females had higher split ratios, reflecting a flexible and risk-spreading strategy. A similar behavior has been described in *Aedes* mosquitoes, and it has been hypothesized that females prioritize skip oviposition in response to varying larval habitat quality in ecologically fluctuating environments such as urban areas (Colton et al., 2003; Harrington & Edman, 2001; Reinbold-Wasson & Reiskind, 2021; Reiter, 2007). Indeed, container-generalists – including *Ae. aegypti* and *Ae. albopictus* – both distribute eggs more evenly among oviposition opportunities, particularly in urban environments with ephemeral containers, whereas *Ae. triseriatus*, which specializes in tree holes, shows reduced skip oviposition (Reinbold-Wasson & Reiskind, 2021). In contrast to *An. coluzzii*, *An. gambiae* females were more conservative and less likely to distribute eggs across oviposition media. This type of behavioral specialization has been reported in another mosquito genus, *Culex*, and may be tailored to adaptation to more hydrologically stable breeding sites in species like *Cx. pipiens* whose larval habitats are semi-permanent and rich in organic content (Allgood & Yee, 2018; Dhileepan, 1997). Oviposition choice studies conducted in *Anopheles* mosquitoes have rarely reported split ratios, and the prevalence of skip-oviposition behavior in the genus remains unknown (Agyapong et al., 2014; Foley & Bryan, 1999; Hardy et al., 2023; Kweka et al., 2011; Lindh et al., 2008; Lowassari et al., 2023; Milugo et al., 2024; Nwaefuna et al., 2019; Osborn et al., 2006; Suh et al., 2016; Sumba et al., 2004).

Among females that allocated eggs to a single oviposition site, the strength of preference also differed between species. *An. gambiae* females, particularly those from field populations, showed highly polarized oviposition preference index (OPI) values, favoring either natal or conspecific field water. This bimodal pattern is consistent with the absence of egg-splitting behavior and underscores a specialized oviposition strategy. In contrast, *An. coluzzii* females displayed more intermediate and widely distributed OPI values, reflecting behavioral plasticity consistent with high skip oviposition. The relationship between ecological specialization and strict oviposition choices has been well documented in herbivorous insects, where specialists show narrow feeding and oviposition preferences while generalists feed and lay eggs on a broad range of host plants (Claridge & Wilson, 1978; Liu et al., 2020; R’Kha et al., 1991; Schäpers et al., 2015; Weeraddana & Evenden, 2022). For example, the specialist, *D. sechellia*, is characterized by high oviposition specificity toward its preferred host, the noni plant, whereas less selective species of the *D. melanogaster* complex typically display a broad range of oviposition preferences (Álvarez-ocaña et al., 2023; Amlou et al., 1998; Dekker et al., 2006). In this study, we have observed OPI values in field-collected *An. gambiae* females that were smaller than those reported for the obligate specialist, *D. sechellia* (Álvarez-ocaña et al., 2023), highlighting the extreme selectivity of some anopheline populations.

The differences in oviposition strategies between *An. gambiae* and *An. coluzzii* may reflect ongoing ecological specialization. Although the cryptic species are broadly sympatric across Africa, they occupy distinct ecological niches at mesogeographic and microspatial scales (Costantini et al., 2009; Diabate et al., 2005; Edillo et al., 2006; Kamdem et al., 2012; Simard et al., 2009). The populations tested here originate from humid regions of Cameroon, where *An. coluzzii* is typically associated with more urbanized environments, whereas *An. gambiae* predominates in rural areas and transient, rain-filled puddles (Kamdem et al., 2012, 2017; Simard et al., 2009; Wondji et al., 2002). A more flexible oviposition strategy in *An. coluzzii* may confer a selective advantage in highly variable aquatic habitats, while *An. gambiae* likely relies on more selective oviposition behavior to optimize offspring survival in ephemeral but hydrologically similar breeding sites. Additionally, larvae of *An. coluzzii* from Yaoundé have higher survival rates than *An. gambiae* in water enriched with ammonium to simulate organic pollution typical of urban habitats, which suggests a link between larval performance and oviposition choice (Tene Fossog et al., 2013).

Further studies are needed to identify the environmental signals that drive oviposition preferences across naturally occurring breeding sites and to assess the performance of immature stages in relation to female choice (Jaenike, 1978). Characterizing the physicochemical composition of breeding waters may help uncover divergent cues and sensory responses between *An. gambiae* and *An. coluzzii*, and reveal the chemical compounds mediating selective oviposition (Lindh et al., 2015). In this study, we did not pursue this approach because of the technical challenges and uncertainty associated with chemically characterizing a diverse set of natural breeding sites sampled over two years. Instead, we combined Pool-seq and individual whole-genome sequencing to assess genetic differentiation and to screen the repertoire of known chemosensory receptor genes. Using this framework, we identified several candidate loci potentially involved in divergent oviposition behaviors between *An. gambiae* and *An. coluzzii*.

### 4.2 Differentiation within the cryptic species is low at small geographic scales

Genome-wide analysis using Pool-seq confirmed that *An. gambiae* and *An. coluzzii* maintain weak but detectable genetic barriers in sympatry, as shown by interspecific *F*_ST_ values ranging from 0.10 to 0.12. Previous studies also reported low genome-wide differentiation among the species pair due the coexistence between extensive hybridization and ecological/genetic divergence (Caputo et al., 2014; Fontaine et al., 2015; Miles et al., 2017). Unsurprisingly, we observed very limited differentiation within each species, with pairwise *F*_ST_ values below 0.010. This homogeneity reflects the absence of reproductive and geographic boundaries expected within the incipient species across the small area where populations were sampled (Cassone et al., 2014; Kamdem et al., 2017).

Genome scans supported this low within-species differentiation, revealing only a few outlier SNPs and no clear genomic signatures of local adaptation at this geographic scale. Unlike previous studies that detected signatures of strong selection at insecticide-associated loci such as target-site resistance or detoxification genes, we found no evidence of such signals within or between species at this geographic scale (Clarkson et al., 2021; Fouet et al., 2018; Grau-bové et al., 2020; Kamdem et al., 2017; Miles et al., 2017). This suggests either reduced variability at xenobiotic resistance loci in our populations or limited resolution of pool-seq to detect fine-scale divergence.

Genome scans thus suggest that divergent oviposition behavior is not underpinned by genomic differentiation between *An. gambiae* and *An. coluzzii*. This finding supports a model in which behavioral divergence can persist between closely related species despite extensive gene flow and genomic homogeneity, a pattern characteristic of members of the *An. gambiae* complex (Coluzzi et al., 1977; Fontaine et al., 2015; Muirhead-Thomson, 1946; Neafsey et al., 2010). Indeed, in the species complex, remarkable phenotypic variations related to host preference, host seeking, and resting behaviors exist within and among species despite limited genomic differentiation due to recent divergence coupled with extensive gene flow (Coluzzi et al., 1977, 1979; Costantini et al., 1999). In such mosaic genomes, instead of large-effect genes that may be detected in whole-genome scans, divergent oviposition behavior may be associated with minor differentiation affecting a subset of chemosensory receptor genes. For example, in closely related species of the *D. melanogaster* complex, high oviposition specificity in *D. sechellia* is modulated by a different odorant-receptor gene tuning compared to less selective species, as well as a contribution of the hexanoic acid receptor Ir75b (Álvarez-ocaña et al., 2023; Auer et al., 2020).

### 4.3 The genetic basis of divergent infochemical perception between species

This study revealed a stark contrast in oviposition preference between *An. gambiae* and *An. coluzzii* that was consistent among field-collected females and long-established colonies maintained in the same environmental conditions. The laboratory strain *An. gambiae* Kisumu strain displayed stricter oviposition preferences than the *An. coluzzii* Ngousso colony, suggesting that the behavioral differences are genetically tractable traits segregating between species. While observed heterozygosity did not vary significantly between species, the mutation load measured by the frequency of heterozygote and homozygote carrying sites across gustatory and ionotropic receptor gene families (Dussex et al., 2023) was higher in *An. gambiae* than in *An. coluzzii*. Detailed analysis of variation at insecticide-associated loci or other gene families point to a relatively higher mutation load in *An. gambiae* compared to *An. coluzzii* due to relaxed purifying selection or distinct demographic histories (Clarkson et al., 2021; Grau-bové et al., 2020; Kamdem et al., 2017; Miles et al., 2017). *F*_ST_ values showed that the level of genetic differentiation between *An. gambiae* and *An. coluzzii* was homogeneous among IRs, GRs, and ORs, suggesting that sensory evolution is driven by divergence acting at the level of individual genes instead of entire families. Within families, there were indeed multiple highly differentiated loci that may be evolving in response to variable chemical stimuli associated with differences in environmental sensing and oviposition behavior between the species.

Candidate outlier genes driving the differentiation of chemosensory receptors between species were AgOr1, AgIr41a, AgIr75g, and AgGr11. A study has shown that knockout of Gr11, which is in the top four genes highly differentiated in the GR family between *An. gambiae* and *An. coluzzii*, leads to longer oviposition site detection time in *Aedes*, suggesting a role for this locus or other gustatory receptor genes in oviposition behavior in mosquitoes (Zhao et al., 2024). Within the IR family, the co-receptor Ir25a was among the least differentiated genes. Co-receptors play a central role in the functioning of tuning receptors and are therefore likely to be under stronger evolutionary constraints, leading to limited differentiation among species (Chen & Amrein, 2017; Sánchez-Alcañiz et al., 2018). Furthermore, poor egg-laying has been reported in *Aedes* Ir25a mutants, consistent with a potential role for this gene in oviposition or reproduction (De Obaldia et al., 2022). In *Drosophila*, response to amine chemostimulants is lost in Ir25a mutants, implying a role in amine compound detection (Abuin et al., 2019). Among tuning receptors, AgIr45a, which is the second most differentiated locus in the IR family has been shown to respond to pyrrolidine and 2-Methyl-2-thiazoline stimuli in *Xenopus* oocyst characterization experiments, which suggests that *An. gambiae* and *An. coluzzii* may diverge in amine detection (Pitts et al., 2017). The most differentiated of all IR loci (AgIr75g) belongs to the AgIR75 clade containing multiple members that mediate sensory responses to carboxylic acids (Cooke et al., 2025; Pitts et al., 2017). The most divergent OR gene identified in this study (AgOr1) is a specialist responding to 4-Methylphenol, a well-known component of human odor plumes used by mosquitoes for long-range attraction, in electrophysiological tests (G. Wang et al., 2010). Although the role of OR genes in responses to oviposition stimulants remains poorly understood, they may contribute to oviposition specialization by shaping variations in long-range detection of volatile compounds between species. This hypothesis is supported by studies demonstrating a causal relationship between differences in tuning properties of Or22a in *D. melanogaster* and *D. sechellia* and species-specific noni attraction (Auer et al., 2020). In addition, in the diamondback moth *Plutella xylostella*, a crucifer specialist, two ORs (Or35 and Or49) are tuned to isothiocyanates, which are secondary metabolites mediating host specialization through divergent oviposition choices (Liu et al., 2020). In summary, highly differentiated loci suggest that *An. gambiae* and *An. coluzzii* diverge in the detection of amines, carboxylic acids, and odorants with potential implications for oviposition specialization. Future functional studies targeting these candidate genes could help elucidate how molecular changes translate into behavioral divergence and ecological specialization between these cryptic species.

## 5. Conclusions

Studies of model organisms such as *D. sechellia* have yielded important insights into specialist–generalist behavioral divergence and have identified key candidate genes underlying oviposition preferences in laboratory settings (Abuin et al., 2019; Álvarez-ocaña et al., 2023; Amlou et al., 1998; Auer et al., 2020; Dekker et al., 2006; McBride, 2007; Reaume & Sokolowski, 2006; R’kha et al., 1997, 1997; Sánchez-Alcañiz et al., 2018). Breakthroughs in non-model systems have similarly revealed the involvement of odorant receptors in oviposition-driven host specialization (Liu et al., 2020), as well as the primary role of divergent visual cues in mediating shifts between specialist and generalist strategies (Y. Wang et al., 2022). Yet despite this progress, we still lack a fundamental understanding of when and how chemosensory receptor genes (and other visual or tactile sensory pathways) diverge between generalists and specialists, or how these sensory signals are integrated into the information-processing mechanisms that drive ecological specialization in natural populations.

Here, we show that *An. gambiae* sensu lato mosquitoes provide a novel system for studying oviposition behavior through specialist and generalist strategies, while minimizing the confounding trade-offs between foraging and oviposition that complicate interpretation in other models such as *Drosophila*. Our findings demonstrate that, at early stages of ecological specialization, behavioral shifts can be mediated by variation at a very small subset of loci against a background of relatively homogeneous genomes. In addition, we found that this small subset of genes is spread across different sensory pathways in the ionotropic, gustatory, and odorant receptor families, underscoring the complexity of chemosensory signaling underlying behavioral differentiation in two cryptic species that only recently split.

However, we acknowledge that the expression profiles of these genes in sensory tissues (antennae, maxillary palps, and other appendages) remain unknown, and functional validation of their roles in oviposition choice was beyond the scope of this study. Future work, including transcriptomics, electrophysiology, and gene-editing approaches, will be essential for determining the precise mechanistic contributions of these loci. Despite these limitations, our study combines behavioral assays, ecological information, and population genomics to provide a framework for investigating the genetics of information processing in behavioral specialists and generalists using a new non-model system that has the potential to advance our understanding of ecological divergence in the wild.

## Acknowledgements

This work was supported by the National Institutes of Health (R01AI150529 to C.K.). We are grateful to the field assistants for their help with larval collections.

## Data availability

Variant call format (VCF) files from the Anopheles gambiae 1000 Genomes Project analyzed in this study are publicly available via MalariaGEN (https://www.malariagen.net/project/ag1000g/). Accession numbers are provided in Supplementary Table S2. Raw Pool-seq sequencing reads generated for this study will be deposited in the NCBI Sequence Read Archive (SRA) under a BioProject accession upon acceptance.

## Author contributions

CF and CK designed the study. MA, CF, FA, AK, SR, DR, CB, VPB and CK performed the research. CF, MA and CK analyzed the data. CK wrote the manuscript. All authors contributed to manuscript revision and approved the final version.

---

**Table S1:** Oviposition preferences of 1,046 gravid *Anopheles* females assessed using two-choice assays.

**Table S2:** List of *Anopheles gambiae* 1000 Genomes Project samples analyzed for diversity and divergence in chemosensory receptor families.

**Table S3:** Genes containing outlier SNPs detected in genome scans of pooled sequencing data. For each gene, genomic coordinates, the number of outlier sites, and the pairwise *F*_ST_ comparisons (between species and/or populations) in which the outliers were detected are provided.

## Notes

### Competing Interest Statement

The authors have declared no competing interest.

